# Reduced gravity and muon flux absence affect *Caenorhabditis elegans* life history traits and viral infection

**DOI:** 10.1101/2024.10.03.616447

**Authors:** Ana Villena-Giménez, Victoria G. Castiglioni, Juan C. Muñoz-Sánchez, Esmeralda G. Legarda, Rubén González, Santiago F. Elena

## Abstract

Environmental conditions fundamentally shape host-pathogen interactions, yet how multiple extreme abiotic stressors combine to influence infection outcomes remains poorly understood. Organisms evolved under specific gravitational and radiation regimes; deviations from these conditions —whether in extreme terrestrial environments or beyond Earth— may alter immune function and pathogen replication. In this study, we investigated the effects of reduced gravity and lowered muon flux on Orsay virus infection in the nematode *Caenorhabditis elegans*. We employed a fully factorial experimental design, examining how each factor, alone and in combination, influences physiological traits and viral load. While below-background radiation radically affected viral accumulation dynamics, reduced gravity had a minor effect. Both factors significantly impacted reproduction and morphology, with some effects magnified by viral infection. These results reveal how even partial modifications of Earth-like gravity and radiation levels can alter pathogen–host interactions. By integrating experimental observations with mathematical modeling, we show that these environmental stressors primarily affect prezygotic reproductive processes and modulate viral replication through distinct and sometimes antagonistic mechanisms. Although this work does not encompass the full complexity of space environments, where cosmic radiation includes high-energy protons and heavy ions, it provides insight into how adjustable models of reduced gravity and radiation can advance our understanding of biological adaptation beyond standard terrestrial conditions.

**IMPORTANCE:** Understanding how extreme environmental conditions affect host–pathogen interactions is critical both for safeguarding biological systems during spaceflight and for exploring fundamental principles of stress biology. This study demonstrates that reduced gravity and diminished muon radiation flux can significantly alter viral infection dynamics and host physiology in *Caenorhabditis elegans*. By integrating experimental data with mathematical modeling, we reveal that these abiotic stresses impact prezygotic reproductive processes and modulate viral replication in distinct and sometimes antagonistic ways. Our findings suggest that even partial deviations from Earth-like conditions can reshape infection outcomes and developmental trajectories, highlighting the need for deeper mechanistic insights into biological adaptation beyond terrestrial norms. These results have implications for space biosciences, evolutionary virology, radiation hormesis theory, and the design of countermeasures to preserve organismal health in extreme or non-terrestrial habitats.

---

Environmental conditions fundamentally shape host-microbe interactions, yet our understanding of how extreme environments alter these relationships remains limited. Organisms evolved under specific gravitational and radiation regimes, and deviations from these conditions may reveal fundamental constraints on physiological and immune function. Such deviations occur in off-Earth environments where organisms may experience a variety of unusual stresses, including reduced gravitational forces (microgravity, μG) and substantial changes in radiation exposure. Understanding how biological systems respond to such perturbations is essential for comprehending life’s responses to conditions beyond Earth: whether encountered during space exploration, in extreme environments, or throughout evolutionary history.

Off-Earth environments can affect the organism’s overall health and immunity. For instance, within relatively short-duration orbital missions (10 - 15 days), blood samples from astronauts showed altered levels of leukocyte distribution and cytokine production and a decrease in the function of Epstein-Barr virus specific CD8^+^ T cells (1). During longer missions (6 months), immune alterations persisted, as evidenced by reduced T-cell function (2), impairment of natural killer cells function (3) and elevated levels of salivary antimicrobial proteins (4). Indeed, the reactivation of latent herpesviruses is well documented in astronauts (4–6): Epstein-Barr virus, varicella-zoster virus and cytomegalovirus infections increased in frequency, duration and amplitude the longer the duration of space missions (5, 7). These observations raise fundamental questions about how gravitational and radiation changes modulate host-pathogen dynamics.

Radiation composition in off-Earth settings differs substantially from that on Earth’s surface. One of the radiation components that differs between space and Earth’s surface is the muon, a charged elementary particle produced when pions decay following cosmic ray interactions with atmospheric molecules. Muons are the main component of cosmic radiation at sea level, with about 600 muons crossing a human body every minute (8). As an important source of radiation for living beings, cells have likely developed mechanisms that buffer muon’s detrimental effects (9, 10). This observation is relevant to radiation hormesis, in which organisms may benefit from low-level radiation exposure but experience detrimental effects when doses deviate significantly in either direction (11). Understanding biological responses to conditions with dramatically reduced muon flux can therefore reveal whether organisms require baseline radiation exposure for optimal function, a key prediction of hormesis theory.

*Caenorhabditis elegans* has long served as a valuable model for studying diverse stresses that can be challenging to dissect in more complex animals. *C. elegans* is a well-characterized metazoan that has been widely employed as a model organism (12). It has many advantages: easy visualization, fast development, a complete characterized cell lineage and genome, and an estimated genetic homology of ∼80% of the protein-coding genome with humans (13). *C. elegans* is exposed to fluctuating environmental conditions in its natural environment, such as heat, cold, food scarcity, oxygen levels, and osmotic perturbations. While repeated mild heat-shock treatments extend *C. elegans* lifespan (14), moderate treatments impair fecundity (15) and embryogenesis (16), and severe treatments decreases longevity (17). These heat-shock phenotypic alterations are synchronized with a strong change in the transcriptional response (18). Exposure to other abiotic stresses, such as hypertonic environments induce transcriptional responses that mimic infection response to a variety of pathogens (19) and heat shocked animals have increased survival upon exposure to *Staphylococcus aureus* (20). In contrast, animals fed with pathogenic bacteria (21) or infected with virus (22) show an increased resistance to heat shock and ubiquitin ligases involved in the intracellular pathogen response (IPR) promote thermotolerance in order to cope with proteotoxic stress, unveiling the close link between biotic and abiotic stresses (23). These findings demonstrate that stress responses and immune pathways are deeply interconnected, raising the question of how non-standard gravitational and radiation environments might modulate infection outcomes.

*C. elegans* has long been used as a model in immunity. It can regulate immune responses recognizing pathogens, influencing the body’s development, and controlling avoidance behavior (24). Although there are significant differences with the immune system of mammals, some mechanisms used to limit pathogenesis show remarkable phylogenetic conservation, such as uridylation and RNAi pathways (25, 26). Since the discovery of Orsay virus (OrV), a lot of effort has been put into the characterization of virus-host interaction dynamics. OrV is a bipartite positive-sense single-stranded RNA virus related to the *Nodaviridae* family that infects *C. elegans* intestinal cells (27). This natural host-pathogen system provides an ideal model for studying how environmental stressors influence real viral infection dynamics, in contrast to artificial infection models. Viral infection produces mild intestinal symptoms such as changes in cytoplasm viscosity, nuclear degeneration and cellular fusion (27), while mild effects in progeny production where only observed in a more susceptible strain (26). Links between viral infection and abiotic stresses have been described in the nematode, with heat stress reducing its susceptibility to OrV (28) and, conversely, OrV-infected animals showing an increased tolerance to heat-shock (22).

The effect of lack of muon radiation flux, hereafter referred as below background radiation (BBR), and of μG on *C. elegans* and other model organisms has been explored before. Van Voorhies *et al*. (29) carried out phenotypic and transcriptomic studies in *C. elegans* at the underground laboratory at the Waste Isolation Pilot Plant (New Mexico, USA). Animals had faster larval growth rates, higher egg laying rates and a different gene expression profile, both at 72 h after being at BBR and after 10 generations. Moreover, a reduction in fertility has been observed in *Drosophila melanogaster* flies reared at the Gran Sasso Underground laboratory (Italy) (30). Studies in simulated μG conditions in *C. elegans* have reported no reproductive and lifespan changes (31), increased intestinal barrier permeability (32) and induction of oxidative stress and dysregulated antioxidant machinery (33).

In this study, we have explored the role of two environmental stressors on the outcome of a viral infection. To do so, we have studied *C. elegans*/OrV pathosystem under a factorial combination of two abiotic factors: gravity intensity (standard and μG), using a Random Positioning Machine (RPM), and muon radiation flux (standard and BBR), achieved by conducting experiments in the Canfranc Underground Laboratory (LSC; Estación de Canfranc, Huesca, Spain). We acclimatized the nematodes to the abiotic stresses and performed the experiments in the third generation of acclimatized animals. This enabled us to investigate long-term effects of μG and BBR in the nematode since it is known that in *C. elegans* responses to stress are transmitted intergenerationally (32). We have measured different fertility traits as a proxy for the animal’s fitness, and body length and morphology to identify developmental impairments. We also assessed intestinal barrier permeability to determine the functional state of the intestine under these stresses, since OrV infects intestinal cells and intestinal collagens are involved in preventing viral entry (34). As a proxy for viral fitness under these different stress conditions, we quantified OrV genomic RNA2 accumulation. Finally, the data was fitted to a fertility model and different parameters associated with prezygotic (early oocyte development, maturation and fertilization rates) and postzygotic (abortion rate) effects estimated. By combining phenotypic observations with mathematical modeling, we reveal how altered gravity and radiation interact to reshape host-pathogen dynamics.

## RESULTS

### The impairment of intestinal barrier by μG is antagonized by BBR

Intestinal integrity is critical for overall health (35), as it is key to nutrient absorption and serves as a primary defense against enteric pathogens, including OrV. Previous work has shown that various abiotic stresses can alter intestinal permeability, with μG causing a significant increase

(31). Accordingly, we sought to validate our simulated microgravity conditions by determining whether they recapitulate this known phenotype. Additionally, we examined how BBR might counterbalance or compound these effects. To measure gut permeability, we employed a dye-based assay in which nematodes are exposed to a blue dye. An intact intestinal barrier restricts the dye to the lumen; conversely, compromised integrity leads to dye leakage into surrounding tissues (Fig. 1A). Firstly, we observed a large and significant overall increase in permeability of animals grown in μG conditions (χ^2^ = 15.882, 1 d.f., *P* < 0.001, 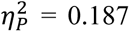), as previously reported. Conversely, BBR conditions were associated to a moderate but still significant reduction in permeability (χ^2^ = 8.387, 1 d.f., *P* = 0.004, 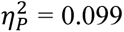). No significant interaction between the two factors has been found (χ^2^ = 0.461, 1 d.f., *P* = 0.497), suggesting that permeability alterations are a consequence of overlapping mechanisms (Fig. 1B). These results validate our experimental system and demonstrate that μG and BBR exert opposing effects on intestinal physiology.

**FIG 1.**
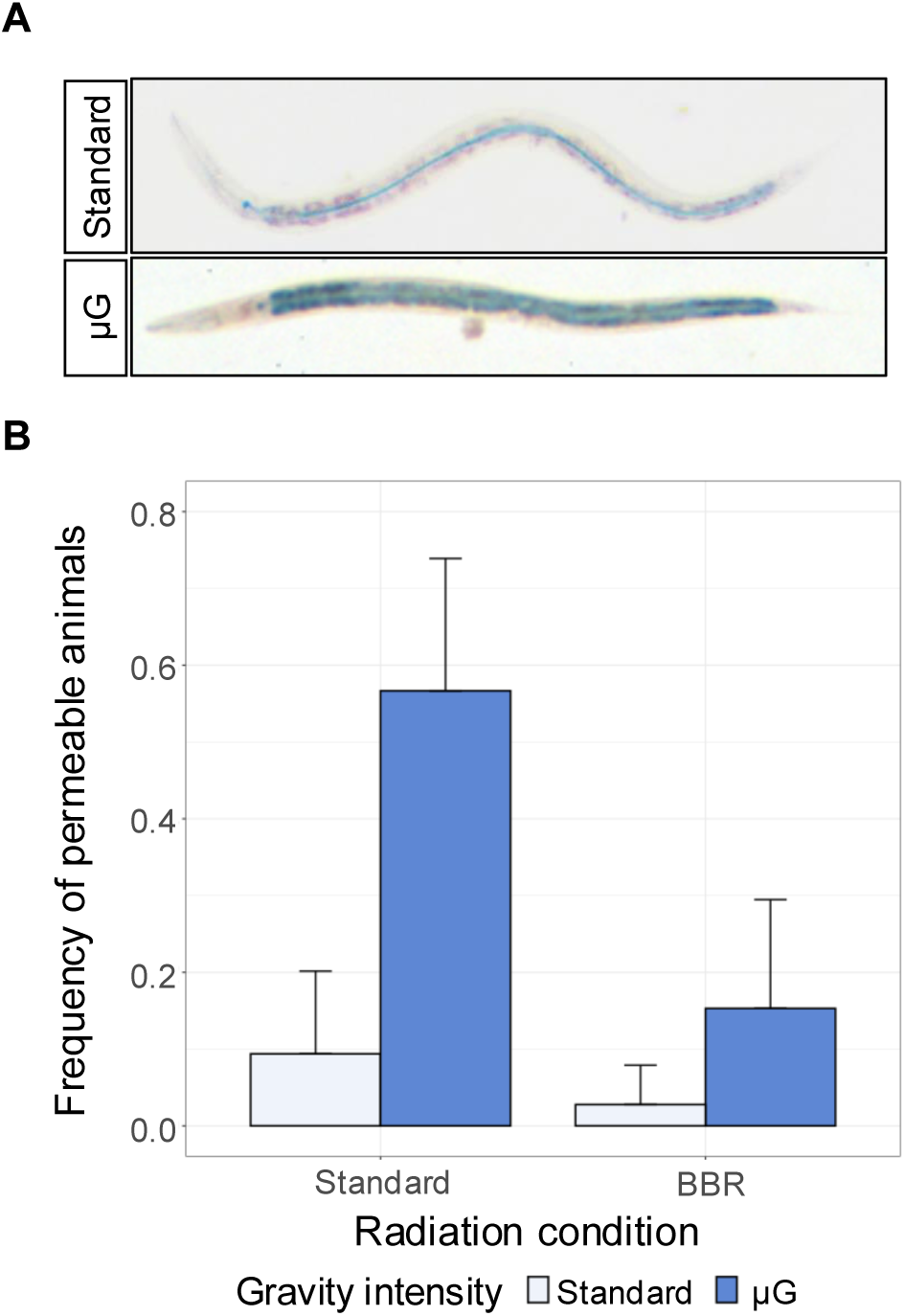
(A) Representative animals in standard (non-permeable) and μG conditions (impaired permeability). Nematodes showing blue staining outside the intestinal tube were scored as permeable whether the salt diffused throughout the body out. (B) Intestinal permeability of animals under standard gravity (*n* = 30), μG (*n* = 28), BBR (*n* = 34) and μG plus BBR (*n* = 24) conditions. Synchronized L1 animals were grown at 20 °C during 48 h and then fed with a solution containing erioglaucine disodium for 3 h. Error bars represent ±1 SEM.

### Viral accumulation dynamics are affected by gravity and radiation intensity

Having confirmed that our simulated conditions produced the expected phenotypic changes in *C. elegans* (*e.g*., increased intestinal permeability under reduced gravity), we next examined whether viral infections remained successful under these modified environments. Our laboratory has previously demonstrated that, under standard laboratory conditions (*i.e*., Earth-like gravity and muon radiation flux), OrV infection in *C. elegans* follows two waves of viral load from L1 larvae to sexually mature adults (36). A first peak is reached approximately 12 hours post-inoculation (hpi) in L1 larvae, which is followed by a sharp decline as antiviral responses are activated by the developing animals, reaching a minimum approximately 18 – 20 hpi (L2 larvae). From there on, a second wider peak is reached that last for at least 18 additional hours (L3 larvae and L4), afterwards, viral load is low, yet viral replication continues at low levels (36). In this work, we observed that under our standard reference conditions, OrV likewise exhibited these two distinct waves, thus confirming the reproducibility of its infection dynamics (Fig. 2, upper left panel). We then asked whether reducing gravity and muon flux would alter either the timing or amplitude of these phases.

**FIG 2.**
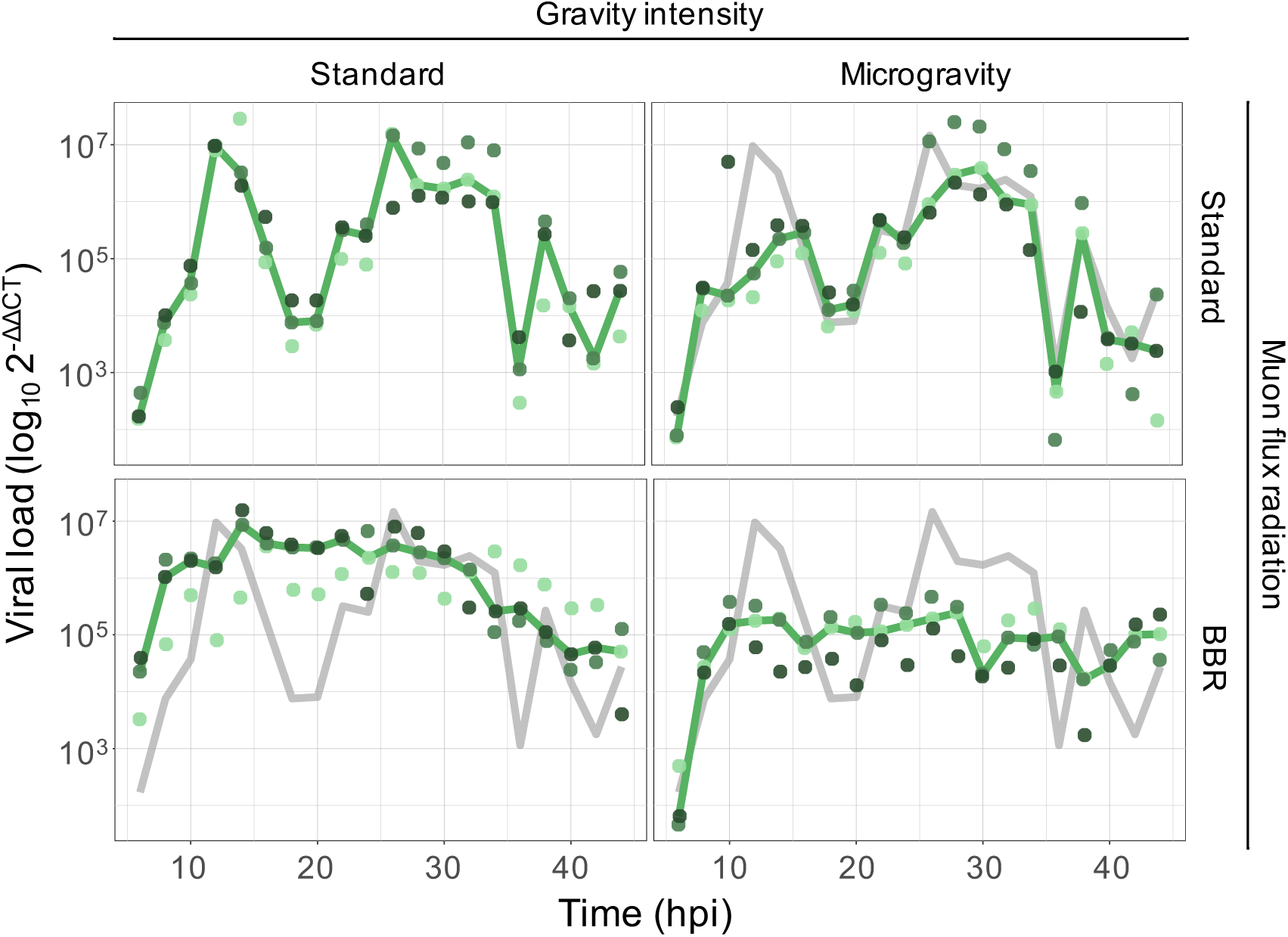
Log-fold change of OrV accumulation, relative to the expression of housekeeping gene *cdc-42*, evaluated in inoculated animals reared under the indicated combination of gravity intensity and muon radiation flux conditions. Different green tones represent independent biological replicates (pools of ∼300 animals per time point). The solid green lines represent the median trajectory. For the sake of comparison, the gray solid lines represent the OrV dynamics observed in standard laboratory conditions (as in the left upper panel).

Gravity intensity has a net significant effect on viral load across time points (χ^2^ = 2255.942, 1 d.f., *P* < 0.001; 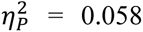). The mean viral load under μG conditions was (5.78 ±0.06)·10^4^,·significantly lower than the (1.02 ±0.01)·10^5^ observed under standard gravity, representing a 1.8-fold reduction over 44 hpi (Fig. 2, upper right panel: green line *vs* gray reference; *post hoc* test, *P* < 0.001). This effect is time-dependent (χ^2^ = 1415.456, 19 d.f., *P* < 0.001; 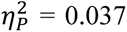), primarily reducing the height of the first peak of viral replication. Muon radiation flux also has a significant impact on viral replication (χ^2^ = 1593.430, 1 d.f., *P* < 0.001; 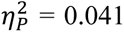). Under BBR conditions, the mean viral load reached (6.36 ±0.07)·10^5^, a 6.3-fold increase compared to normal muon flux (Fig. 2, lower left panel: green line *vs* gray reference). The effect of BBR strongly altered the temporal dynamics of OrV accumulation (χ^2^ = 3103.138, 19 d.f., *P* < 0.001; 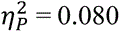), eliminating the biphasic replication pattern: a single peak emerges around 12 hpi, remains stable until 36 hpi, and then gradually declines.

These environmental effects were not independent. A significant interaction was detected between gravity and radiation conditions (χ^2^ = 1636.075, 1 d.f., *P* < 0.001; 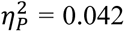), which was also time-dependent (χ^2^ = 1934.595, 19 d.f., *P* < 0.001; 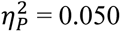). The combined stress of μG and BBR produces a distinctive pattern resembling the BBR conditions but with a reduction in viral load similar to the one observed in μG —approximately 1.8-fold lower than the control throughout the entire developmental period (Fig. 2, lower-right panel: green line *vs* gray reference; *post hoc* test, *P* < 0.001).

In summary, μG exerts a negative effect on viral replication, BBR enhances it while altering infection dynamics, and their combination results in a compounded reduction of viral accumulation.

### Fecundity traits are affected by reduced gravity and radiation

OrV infections are typically described as having a mild impact on *C. elegans* fitness (37). To determine whether this impact might be altered by our environmental treatments, we performed time-series measurements of three fecundity-related traits —number of viable progeny (*A*), number of unfertilized eggs (*U*), and number of nonviable embryos (*N*)— in four conditions: standard environment, μG, BBR, and the combination of μG plus BBR (Fig. S1). These measurements were done both in non-inoculated animals (control; blue lines and circles) and in OrV-inoculated ones (OrV; green lines and triangles). All these data were fitted to the generalized linear mixed model (GLMM) described in the Materials and Methods section. The results from these analyses are shown in Table 1. For a detailed description of the effect of the different stresses on *A*, *U* and *N*, we refer to the supplementary text in Supplementary Materials online. Here, we will focus on the combination of these three traits into a more eco-evolutionary relevant trait, the per-day reproductive success defined as *R*(*t*) = *A*(*t*)/[*A*(*t*) + *N*(*t*) + *U*(*t*)] (Fig. 3). Focusing first on the effects of the environmental conditions in non-inoculated animals, we found that differences in gravity intensity had a significant effect, though of small magnitude (Table 1; *P* < 0.001, 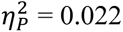), with animals reared at standard gravity having an average *R* of 0.950 ±0.005 that was reduced to 0.927 ±0.004 in μG (Fig. 3, upper left *vs* upper right panels, blue). The effect of muon radiation flux was conditional to the value of gravity intensity (Table 1; *P* = 0.002, 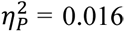). In μG, *R* was not affected by differences in muon radiation flux (*post hoc* test, *P* = 0.691) (Fig. 3, upper left *vs* lower right panels, blue). However, in standard gravity conditions the minimization of muon radiation flux was associated with a reduction in average *R* from 0.962 ±0.007 to 0.939 ±0.006 (*post hoc* test, *P* < 0.001) (Fig. 3, upper left *vs* lower left panels, blue).

**FIG 3.**
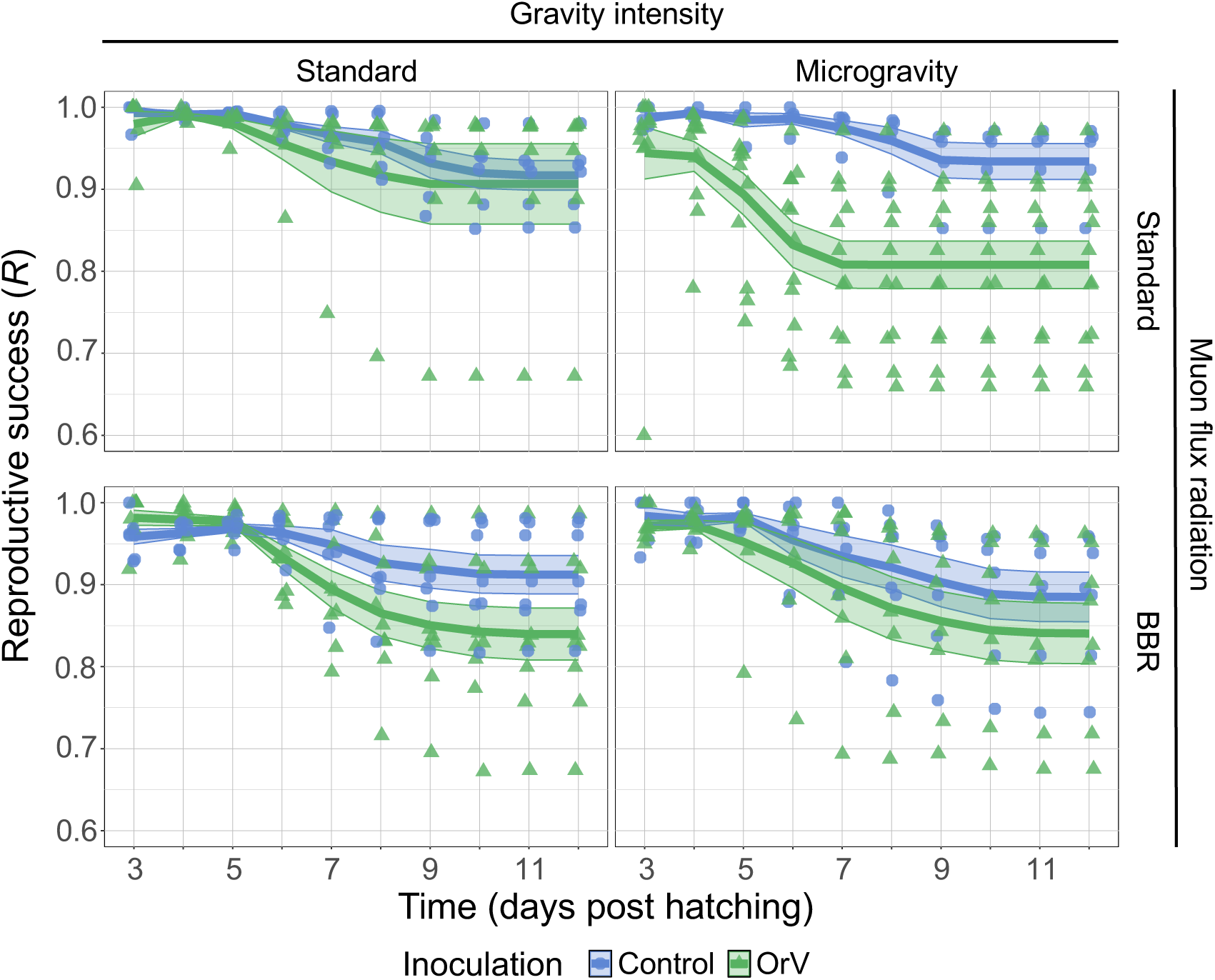
Per day reproductive success (*R*) of OrV-inoculated (green) and non-inoculated (blue) animals under different combinations of environmental stresses. Color triangles and circles represent replicates and solid lines the mean of replicates with ±1 SEM plotted as a ribbon. Number of non-inoculated animals: *n* = 6 for standard conditions, *n* = 5 for μG, *n* = 7 for BBR, and *n* = 7 for μG and BBR. Number of inoculated animals: *n* = 6 for standard conditions, *n* = 12 for μG, *n* = 9 for BBR, and *n* = 8 for BBR plus μG.

**TABLE 1.** Results of the fecundity data fitting to the GLMM testing for the effect of gravity intensity, muon radiation flux and OrV infection. Significant effects of small magnitude are highlighted in light gray, while significant effects of medium and large magnitude are highlighted in dark gray.

| Source of variation | d.f. | Unfertilized eggs (*U*) |  |  | Nonviable embryos (*N*) |  |  | Progeny (*A*) |  |  | Reproductive success (*R*) |  |  |
| --- | --- | --- | --- | --- | --- | --- | --- | --- | --- | --- | --- | --- | --- |
| | | *F* | *P* | $\eta_P^2$ | *F* | *P* | $\eta_P^2$ | *F* | *P* | $\eta_P^2$ | *F* | *P* | $\eta_P^2$ |
| Model | 7, 592 | 3.113 | 0.003 | 0.036 | 11.702 | < 0.001 | 0.122 | 13.129 | < 0.001 | 0.134 | 19.737 | < 0.001 | 0.189 |
| Gravity intensity | 1, 592 | 1.923 | 0.166 | 0.003 | 3.825 | 0.051 | 0.006 | 22.538 | < 0.001 | 0.037 | 13.221 | < 0.001 | 0.022 |
| Muon radiation flux | 1, 592 | 1.122 | 0.290 | 0.002 | 5.010 | 0.026 | 0.008 | 0.410 | 0.522 | 0.001 | 0.425 | 0.515 | 0.001 |
| Infection status | 1, 592 | 3.864 | 0.050 | 0.006 | 9.244 | 0.002 | 0.015 | 19.131 | < 0.001 | 0.031 | 47.457 | < 0.001 | 0.074 |
| Gravity × Radiation | 1, 592 | 1.866 | 0.172 | 0.003 | 23.054 | < 0.001 | 0.037 | 21.024 | < 0.001 | 0.034 | 9.599 | 0.002 | 0.016 |
| Gravity × Infection | 1, 592 | 1.968 | 0.161 | 0.003 | 12.585 | < 0.001 | 0.021 | 13.209 | < 0.001 | 0.022 | 14.810 | < 0.001 | 0.024 |
| Radiation × Infection | 1, 592 | 1.624 | 0.203 | 0.003 | 15.129 | < 0.001 | 0.025 | 1.097 | 0.295 | 0.002 | 8.321 | 0.004 | 0.014 |
| Gravity × Radiation × Infection | 1, 592 | 2.020 | 0.156 | 0.003 | 0.259 | 0.611 | 0.000 | 6.689 | 0.010 | 0.011 | 10.964 | < 0.001 | 0.018 |

*R* was also negatively affected by infection itself (Table 1; *P* < 0.001, 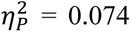), with inoculated animals having grand mean *R* of 0.917 ±0.004 and non-inoculated animals of 0.961 ±0.005, an effect of moderate magnitude. The effect of inoculation was mediated by gravity intensity (Table 1; *P* < 0.001, 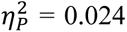). The mean *R* of inoculated animals exposed to μG (0.894 ±0.005) was significantly lower than in standard gravity (0.941 ±0.006; *post hoc* test, *P* < 0.001) (Fig. 3, upper left *vs* upper right panels, difference between blue and green). The effect of inoculation was also mediated by the muon radiation flux (Table 1; *P* = 0.004, 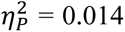). In this case, *R* of inoculated animals in BBR conditions (0.910 ±0.006) was lower than for animals inoculated in standard muon radiation flux (0.949 ±0.006; *post hoc* test, *P* = 0.020) (Fig. 3, upper left *vs* lower left panels, difference between blue and green). Finally, the effect of inoculation significantly depended on the interaction between the two environmental conditions (Table 1; *P* < 0.001, 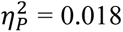). The largest effect on *R* was observed for inoculated animals grown under μG and standard muon radiation flux (0.866 ±0.007) (Fig. 3, upper left *vs* lower right panels, difference between blue and green), while the smallest effect was seen for non-inoculated animals under the same combination of factors (0.97 ±0.01) (green, upper left *vs* lower right panels in Fig. 3).

In conclusion, both environmental stressors impacted reproductive success, though their effects varied across traits. μG always reduced reproductive success. BBR reduced reproductive success in standard gravity, and its effects were smaller or absent in μG. In standard conditions, OrV negatively impacted the animals’ reproductive success, especially by increasing the number of unfertilized eggs. These effects are worsened by the combination of μG and BBR, indicating that these stresses likely affect different pathways of the infection response.

### The observed effects of BBR and viral infection are likely prezygotic

The above statistical descriptions and analyses highlight the effect of the three experimental factors in the data. However, this sort of analyses fails to provide a mechanistic explanation for the observed differences. To dig into possible mechanisms, we developed a simple model that describes the processes from the maturation of germline precursors up to the hatching of eggs to liberate L1 larvae. The model is described by the set of six ordinary differential equations given by Eqs. 1 – 6 in Materials and Methods. The model includes five parameters to describe different processes. Fitting the data shown in Fig. 1 into Eqs. 1 – 6 allows to disentangle whether observed differences in *A*, *N* and *U* under different environments and infection status can be attributable to pre- or post-zygotic effects. Table 2 shows the fitting estimates for each condition (see Fig. S2 for illustration of the best fittings). Table S1 shows the results of the three-ways ANOVA model fitted to the parameter estimates. Gravity intensity itself had no significant effect on any of the six model parameters. Muon radiation flux had significant effects, of large magnitude, in the rates of germline precursors differentiation into mature oocytes (*δ*) as well as in the rate of oocyte migration from the gonads into the spermathecae (*η*). Animals reared in BBR conditions experienced a reduction in both parameters, in the case of *δ* from 0.65 ±0.03 d^−1^ to 0.45 ±0.02 d^−1^ (*post hoc* test *P* < 0.001); in the case of *η* from 0.92 ±0.06 d^−1^ to 0.69 ±0.06 d^−1^ (*post hoc* test *P* = 0.008).

**TABLE 2.**
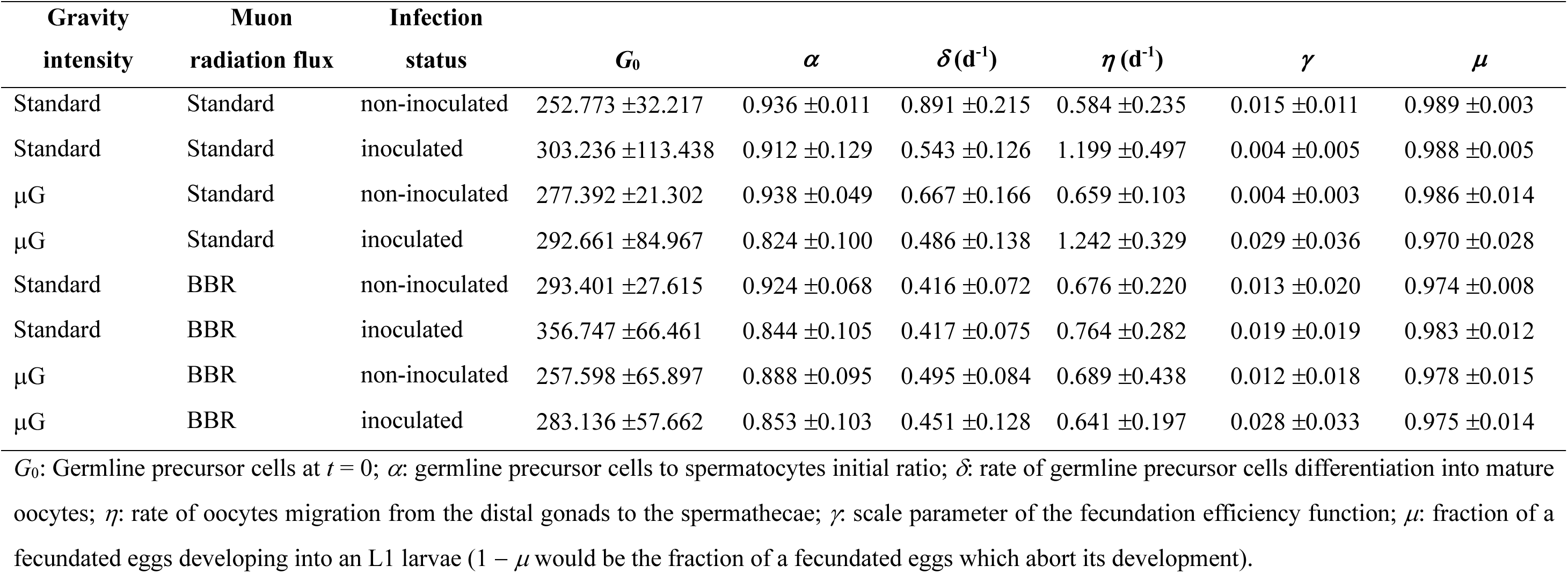
Estimation of model parameters and initial condition. Values represent the mean of the replicates (±1 SD). See Fig. S2 for the fitting to the experimental data.

| Gravity<br>intensity | Muon<br>radiation flux | Infection<br>status | $G_0$ | $\alpha$ | $\delta(\text{d}^{-1})$ | $\eta(\text{d}^{-1})$ | $\gamma$ | $\mu$ |
| --- | --- | --- | --- | --- | --- | --- | --- | --- |
| Standard | Standard | non-inoculated | 252.773 ± 32.217 | 0.936 ± 0.011 | 0.891 ± 0.215 | 0.584 ± 0.235 | 0.015 ± 0.011 | 0.989 ± 0.003 |
| Standard | Standard | inoculated | 303.236 ± 113.438 | 0.912 ± 0.129 | 0.543 ± 0.126 | 1.199 ± 0.497 | 0.004 ± 0.005 | 0.988 ± 0.005 |
| $\mu\text{G}$ | Standard | non-inoculated | 277.392 ± 21.302 | 0.938 ± 0.049 | 0.667 ± 0.166 | 0.659 ± 0.103 | 0.004 ± 0.003 | 0.986 ± 0.014 |
| $\mu\text{G}$ | Standard | inoculated | 292.661 ± 84.967 | 0.824 ± 0.100 | 0.486 ± 0.138 | 1.242 ± 0.329 | 0.029 ± 0.036 | 0.970 ± 0.028 |
| Standard | BBR | non-inoculated | 293.401 ± 27.615 | 0.924 ± 0.068 | 0.416 ± 0.072 | 0.676 ± 0.220 | 0.013 ± 0.020 | 0.974 ± 0.008 |
| Standard | BBR | inoculated | 356.747 ± 66.461 | 0.844 ± 0.105 | 0.417 ± 0.075 | 0.764 ± 0.282 | 0.019 ± 0.019 | 0.983 ± 0.012 |
| $\mu\text{G}$ | BBR | non-inoculated | 257.598 ± 65.897 | 0.888 ± 0.095 | 0.495 ± 0.084 | 0.689 ± 0.438 | 0.012 ± 0.018 | 0.978 ± 0.015 |
| $\mu\text{G}$ | BBR | inoculated | 283.136 ± 57.662 | 0.853 ± 0.103 | 0.451 ± 0.128 | 0.641 ± 0.197 | 0.028 ± 0.033 | 0.975 ± 0.014 |

$G_0$ : Germline precursor cells at $t = 0$ ; $\alpha$ : germline precursor cells to spermatocytes initial ratio; $\delta$ : rate of germline precursor cells differentiation into mature oocytes; $\eta$ : rate of oocytes migration from the distal gonads to the spermathecae; $\gamma$ : scale parameter of the fecundation efficiency function; $\mu$ : fraction of a fecundated eggs developing into an L1 larvae ( $1 - \mu$ would be the fraction of a fecundated eggs which abort its development).

Infection status had a significant effect of large magnitude in *δ* and *η* (Table S1; *P* < 0.001, 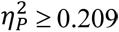 in both cases) and significant but of moderate magnitude in the initial number of germline precursors (*G*_0_) and in spermatocyte to germline precursors initial ratio (*α*) (Table S1; *P* ≤ 0.038, 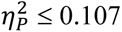 in both cases). Inoculated animals show a reduction in *δ* from 0.62 ±0.03 d^−1^ to 0.47 ±0.02 d^−1^ (*post hoc* test *P* < 0.001), an increase in *η* from 0.65 ±0.06 d^−1^ to 0.96 ±0.06 d^−1^ (*post hoc* test *P* < 0.001), a reduction in *α* from 0.92 ±0.02 to 0.86 ±0.02 (*post hoc* test *P* = 0.016), and an increase in *G*_0_ from 270 ±14 to 309 ±12 (*post hoc* test *P* = 0.038).

Table S1 also shows that some of the effects involved interactions between the three environmental factors. Firstly, *δ* was affected by gravity intensity in a manner dependent on the muon radiation flux (significant interaction gravity by radiation term; *P* = 0.006, 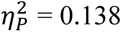: in standard muon radiation flux and standard gravity *δ* = 0.72 ±0.04 d^−1^, but reduced to *δ* = 0.58 ±0.03 d^−1^ in μG (19.7% reduction; *post hoc* test *P* = 0.003); in BBR conditions and standard gravity *δ* = 0.42 ±0.03 h^−1^, statistically undistinguishable from *δ* = 0.47 ±0.03 d^−1^ in μG (*post hoc* test *P* = 0.107). Secondly, the magnitude of the effect of muon radiation flux on *δ* and *η* also depended on the infection status (Table S1; *P* ≤ 0.001, 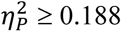 in both cases). In the case of *δ*, non-inoculated animals reared in standard muon radiation flux conditions had *δ* = 0.78 ±0.04 d^−1^, while at BBR conditions, *δ* = 0.46 ±0.03 d^−1^, representing a significant reduction (*post hoc* test *P* < 0.001); inoculated animals grown at standard muon radiation flux conditions had *δ* = 0.51 ±0.03 d^−1^, while at BBR conditions, *δ* = 0.43 ±0.03 d^−1^, representing a significant reduction (*post hoc* test *P* = 0.036). In other words, the magnitude of the effect on *δ* due to muon radiation flux was larger in non-inoculated animals than in inoculated ones (41.6% *vs*. 15.56% reductions). In the case of *η*, non-inoculated animals reared in standard muon radiation flux had *η* = 0.6 ±0.1 d^−1^, while at BBR conditions, *η* = 0.68 ±0.08 d^−1^, values that were not significantly different (*post hoc* test *P* = 0.315); inoculated animals grown at standard muon radiation flux conditions had *η* = 1.22 ±0.08 d^−1^, while at BBR conditions, *η* = 0.70 ±0.08 d^−1^, representing a significant reduction (*post hoc* test *P* < 0.001). In other words, the magnitude of the effect on *η* due to muon radiation flux was larger in inoculated animals than in non-inoculated ones (44.0% reduction *vs*. 9.8% nonsignificant increase).

In conclusion, we found that the three environmental factors had significant effects in parameters related to prezygotic processes, with the magnitude and sign of the observed changes depending on the precise combination of factors. In terms of wider effects, OrV infection affected more developmental processes than either muon radiation flux or gravity intensity.

### Synergistic effects of μG, BBR and infection in developmental rates

To better understand the effects of these environmental factors in another key developmental/health metric, we next focused on larval development (Fig. 4A). There was a significant overall effect of gravitational differences yet of small magnitude (χ^2^ = 45.734, 1 d.f., *P* < 0.001, 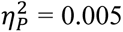), with animals raised in μG being larger than those raised under standard conditions (mean values across time points: 678 ±4 µm *vs* 640 ±4 µm) (Fig. 4A, upper left *vs* upper right panels, blue). Furthermore, the difference is size increased with time post-hatching (χ^2^ = 48.942, 3 d.f., *P* < 0.001; 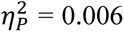). Muon radiation flux also had a significant overall effect of small magnitude (χ^2^ = 28.179, 1 d.f., *P* < 0.001, 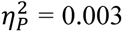), with nematodes exposed to BBR conditions being larger than those in standard conditions (674 ±4 µm *vs* 644 ±4 µm) (Fig. 4A, upper left *vs* lower right panels, blue). The effect magnitude was also time-dependent (χ^2^ = 17.747, 3 d.f., *P* < 0.001; 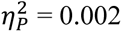). Inoculation had a significant impact as well (χ^2^ = 20.630, 1 d.f., *P* < 0.001, 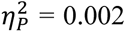), with inoculated animals being smaller than their non-inoculated counterparts (647 ±4 µm *vs* 672 ±4 µm). The effect magnitude varied over time, generally increasing (χ^2^ = 14.983, 3 d.f., *P* = 0.002; 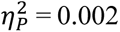), though not uniformly across all conditions.

**FIG 4.**
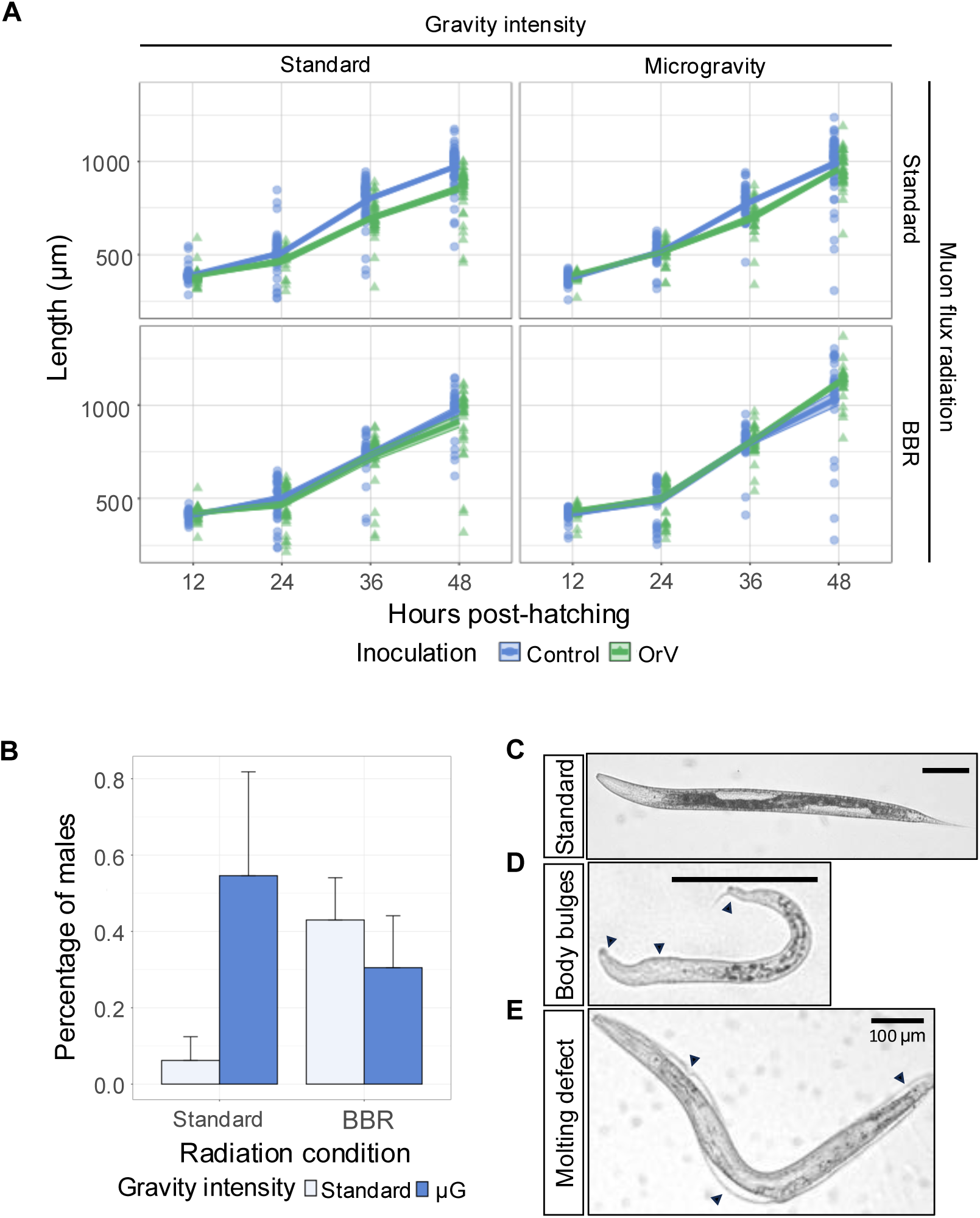
Growth and development of non-inoculated and OrV-inoculated animals under different gravity and radiation conditions. (A) Starting from plates with 400 individuals, photographs were taken at 12, 24, 36, and 48 hours after hatching to assess the effect of the stress conditions in larval development. Solid lines represent the mean and ribbons represent ±1 SEM (number of replicates ranged from 19 to 130). (B) Percentage of males quantified 48 h after hatching under standard (*n* = 1606), μG (*n* = 733), BBR (*n* = 3490), and BBR plus μG (*n* = 1640). Error bars represent ±1 SEM. (C) Animal grown in standard conditions. (D) Individual with bulges around the body and short length. Extracted from the population of 24 h in BBR conditions; scale bar of 100 μm. (E) Individual defective in molting. Extracted from the population 48 h in BBR plus μG; scale bar of 100 μm.

The magnitude of these effects depended on the pairwise interactions of the three factors (χ^2^ ≥ 10.424, 1 d.f., *P* ≤ 0.001, 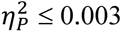 in all combinations) and varied over time (χ^2^ ≥ 12.269, 3 d.f., *P* ≤ 0.007, 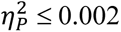 in all combinations). Focusing first in the interaction between gravity intensity and muon radiation flux, animals grown in μG under BBR conditions were, on average, larger than those grown only in μG (702 ±6 µm *vs* 654 ±5 µm; *post hoc* test, *P* < 0.001) (Fig. 4A, upper right *vs* lower right panels, blue) and larger than those grown only under BBR conditions (702 ±6 µm *vs* 646 ±6 µm; *post hoc* test, *P* < 0.001) (Fig. 4A, lower left *vs* lower right panels, blue lines). Now looking at the interaction between gravity intensity and inoculation, we found that under standard gravity intensity, inoculated animals were on average smaller than non-inoculated ones (618 ±6 µm *vs* 671 ±4 µm; *post hoc* test, *P* < 0.001) (Fig. 4A, upper left panel, differences between blue and green), while inoculation had no significant effect on growth under μG conditions (675 ±6 µm *vs* 673 ±6 µm; *post hoc* test, *P* = 0.602) (Fig. 4A, upper right panel, differences between blue and green). Finally, exploring the interaction between muon radiation flux and inoculation, we observed that inoculation had no net effect on animals grown under BBR conditions (675 ±6 µm *vs* 673 ±6 µm; *post hoc* test, *P* = 1) (Fig. 4A, lower left panel, differences between blue and green). Three-way interactions were not statistically significant either independently (χ^2^ = 0.225, 1 d.f., *P* = 0.636) or over developmental time (χ^2^ = 2.034, 3 d.f., *P* = 0.565), indicating that the effect of infection was independent on the interaction between the two abiotic stresses (Fig. 4A, lower right panel, differences between blue and green).

Populations of *C. elegans* are typically dominated by hermaphrodites, with males arising infrequently under standard conditions, only once per 1606 individuals in our experiments. Although rare, males can be biologically significant by enabling outcrossing, which increases genetic diversity and may facilitate adaptation under stressful conditions (38). Notably, the frequency of males increased when nematodes were subjected to reduced gravity or BBR. For instance, we found four males in a population of 733 under μG, 15 out of 3490 under BBR alone, and five out of 1640 under combined μG and BBR (Fig. 4B). While neither gravity intensity (χ^2^ = 2.606, 1 d.f., *P* = 0.106) nor muon ration flux (χ^2^ = 1.431, 1 d.f., *P* = 0.232) alone significantly affected the number of males produced, their combination led to a ∼5-fold increase in male frequency compared to standard conditions, a statistically significant yet small effect (χ^2^ = 5.315, 1 d.f., *P* = 0.021, 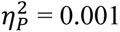).

In addition, under BBR conditions we observed dumpy animals, characterized by a shorter and stouter body morphology due to defects in structural genes like collagens (3 in a population of 1843 individuals), as well as other phenotypes that deviate from the standard (Fig. 4C), which included increased head-length proportion, body bulges (Fig. 4D) and molting defects (Fig. 4E). Neither of these phenotypes were observed in standard conditions (Binomial test, *P* < 0.001). A possible explanation is that lack of radiation might affect genes that are involved in molting, such as collagens (39).

In conclusion, OrV inoculation mitigates the effect of the two abiotic stresses on developmental rates, the two abiotic stresses result in increased male frequency and BBR also causes diverse morphological abnormalities.

## DISCUSSION

In this study, we sought to investigate the independent and combined effects of two environmental stressors (reduced gravity and absence of muon radiation flux) on a biological system comprising *C. elegans* and its natural viral pathogen OrV. Organisms evolved under specific gravitational and radiation conditions at Earth’s surface; deviations from these parameters provide opportunities to understand the constraints these factors impose on physiology and immunity. Previous work suggested that even partial modifications of Earth-like gravity and radiation levels could influence immune responses and pathogen–host interactions (7, 40, 41). Consequently, we focused primarily on reduced muon flux (the major component of cosmic radiation at sea level) in an underground laboratory, and on simulated reduced gravity using an RPM. Although space environments include additional stressors such as heavy-ion radiation and solar particle events (42), our results demonstrate how extreme environments can fundamentally alter host-pathogen interactions.

We first validated that our reduced-gravity simulation recapitulated previously reported phenotypes, such as increased intestinal permeability under microgravity (32). Although agar plate-based RPM experiments do not fully mimic true weightlessness, and surface tension can interfere with microgravity simulation fidelity (43), we still observed a significant compromise in intestinal integrity under μG. Furthermore, RPM microgravity simulation in agar plates has been proved to induce changes in *C. elegans* physiology (31), while liquid culture in a RPM produces a shear stress that should be acknowledged (44). It is also worth-mentioning that liquid culture in *C. elegans* has an impact in gene expression (45) and growth (46), which represents a disadvantage when considering other microgravity simulation systems. Conversely, culturing nematodes under BBR (*i.e*., drastically reduced muon flux) partially offset μG-induced intestinal permeability, but their combination worsens reproductive success, suggesting that microgravity and BBR elicit overlapping but distinct physiological responses.

Our results confirm that μG and BBR represent stressful environments that affect responses to viral infections and development (1–7, 29). Regarding development and fecundity, the small but significant increase in male frequency in the combination of the two stresses suggests that off-Earth conditions may disrupt cellular homeostasis, potentially affecting chromosomal disjunction as observed with other environmental stresses (47). Increased male frequency could have major population-level consequences, as increased mating increases genetic diversity through allele shuffling, bolstering adaptation to stressful conditions (38, 48). Animals grown under μG or BBR conditions were larger, with their combination producing further increases. These μG results are in disagreement with previous works, which reported decreased body length in simulated microgravity (33) and in spaceflight (49), likely due to our experimental design: animals were preadapted to μG during two generations, grew on solid medium, and experienced different exposure timing and duration without additional stresses like bleaching synchronization. Regarding viral infection, OrV-inoculated animals in standard conditions were shorter than non-inoculated, while it had no effect under other stresses. This confirms early observations of OrV infection effects on development (50) and consistent with observations in adult nematodes expressing human hepatitis delta viral antigens (51). Regarding viral infection dynamics, μG had a small effect in the shape of the virus accumulation curve, reducing the accumulated viral load over 44 hpi. BBR had major effects in OrV accumulation dynamics, erasing the distinction between both viral production phases and resulting in a large reduction in accumulated OrV. Combining both stresses fully changed temporal viral production shape, resembling BBR patterns but further reducing accumulated viral load. These observations suggest that (*i*) simulated μG conditions may cause minor nematode immune system impairment and (*ii*) BBR-induced stress on nucleic acids stability impacts OrV replication through unknown mechanisms. Our results are consistent with spaceflight-induced immune alterations and herpesvirus reactivation in astronauts (3, 5–7, 52).

The molecular mechanisms underlying these effects likely involve multiple conserved pathways. Short-term exposures to μG cause small, reversible *C. elegans* transcriptomic alterations, while prolonged expositions result in larger transgenerational reversible effects (31, 53, 54). These alterations affect longevity-regulating insulin/IGF-1 (55) and sphingolipid signaling pathways (56), neuronal function and/or cellular metabolism mediated by the *daf-16*/FOXO signaling pathway (57), and misregulation of responses to oxidative stress and the antioxidant defense system (33). Some of these pathways are also linked to OrV infection (55), which generates reactive oxygen species (ROS) as part of the immune response, creating oxidative stress (56). Failure to produce ROS antiviral responses would result in enhanced virus accumulation. Our fertility modeling approach revealed effects appear largely attributable to differences in prezygotic parameters, such as oocyte maturation and fertilization rates. This aligns with findings showing altered major sperm protein genes (MSP) regulation in BBR condition (29). MSPs are involved in sperm motility and amoeboid-like crawling (57) and are critical for oocyte and egg development, ovulation, spermathecal valve dilation, and parthenogenesis (58, 59). Given MSP functional diversity, altered *msp* gene expression under BBR conditions likely reflects in our model’s prezygotic parameters.

Our study is amongst the first exploring biological implications of muon radiation flux absence on viral infection. While most research focused on cosmic radiation exposure on natural background radiation, the reduction of secondary cosmic rays, including muons, presents an underexplored variable (29). No study has explored radiation absence effects on viral infection progression, despite potential contributions to viral activation in astronauts. Muon radiation has been present during organisms’ evolution as a result organisms have developed mechanisms directed to cope with the effects of this type of radiation in the integrity of their genetic material integrity and physiological processes. The negative impact of lack of muon radiation in reproductive success aligns with the emerging theory of radiation as an hormetic agent (11), where doses below and above natural background may harm organisms.

Combined μG and BBR impact creates a singular environment affecting viral infection dynamics. These off-Earth stresses are not isolated but interact to modulate viral reactivation or replication. The immune dysregulation caused by μG, when compounded by the altered radiation environment, demonstrates how environmental factors integrate to produce emergent biological properties not predictable from individual treatments. The systems-level interactions we observed (where reduced gravity and radiation absence create overlapping yet distinct physiological responses) highlight the complex dependencies organisms have evolved under Earth’s environmental conditions.

In conclusion, evidence presented here highlights the intricate interplay between reduced gravity, diminished muon flux and viral infection. Even introducing a couple of off-Earth conditions (microgravity and below-background radiation) substantially reshape host physiology and pathogen replication. Our findings suggest that lowered gravity may impair certain *C. elegans* immune function aspects, allowing modest viral load increases, while the lack of muon flux alters viral accumulation dynamics more pronouncedly. These stressors create an environment in which viral activity and host fitness are influenced by overlapping yet distinct mechanisms. For organisms evolved in Earth’s gravitational field and background radiation, deviations in these parameters (whether in orbital stations, underground conditions, or on other planetary bodies) can induce notable biological effects. Further research involving a wider range of radiation types and additional parameters will help clarify the mechanisms behind these observations and guide strategies to preserve health under non-terrestrial conditions.

## MATERIAL AND METHODS

### *C. elegans* strains and culturing

*C. elegans* was cultured and maintained at 20 °C on nematode growth media (NGM) agar plates seeded with *Escherichia coli* OP50. ERT54 (*jyIs8[pals-5p::GFP + myo-2p::mCherry]X*), a transgenic strain with a genetic wild-type (Bristol N2) background that expresses GFP in response to intracellular infection (60) was used for all experiments. The more susceptible strain SFE2 (*drh-1[ok3495]IV;mjls228*) was used to produce OrV stocks.

In order to obtain synchronized animal populations, plates with embryos were carefully washed with M9 buffer to remove larvae and adults but leaving the embryos behind. Plates were washed again using M9 buffer after 1 h to collect larvae hatched within that time span and transferred to seeded NGM plates.

A variable number of plates of 500 synchronized L1 animals were used to assess male frequency. Plates were stored at 20 °C and, two days after hatching, males and total population were counted using a Zeiss Stemi305 stereomicroscope.

### Experimental design and locations

Four different environmental conditions have been tested in this work: standard conditions, μG, BBR, and the combination of μG plus BBR. Experiments conducted in standard and microgravity conditions were performed in the Evolutionary Systems Virology laboratory in the Institute for Integrative Systems Biology (Paterna, Spain). The experiments that included BBR conditions were carried out in the LSC in three different periods from 2022 to 2024 due to technical issues.

μG was simulated using an RPM (Yuri Gravity GmbH) set to the zero-gravity mode. Values of G-force were monitored through the RPM software and maintained at approximately 0.001 − 0.002 g throughout the experiments. The RPM was placed inside an incubator at 20 °C. Animals used in experiments of μG conditions were acclimated along two generations, with studies done on the third generation and its progeny. Plates were sealed with parafilm to maintain their humidity.

BBR experiments were performed in the LSC LAB2400. The measured integrated muon radiation flux in this facility is ∼5×10 ^−3^ m^−2^ s^−1^ (61). For comparison, the muon radiation flux at sea level in the Northern hemisphere is about 150 m^−2^ s^−1^ (62). Thermoluminescent dosimeters sensitive to various sources of radiation are placed at different locations of the underground laboratory. In 2022 they showed and average dose rate of 0.71 ±0.03 mSv per year in the underground facilities, compared with the 1.36 ±0.03 mSv per year of the above ground laboratory (63). Radon radiation from the mountain was reduced to 1 mBq m^−3^ thanks to a good ventilation system and a Radon Abatement System, a value substantially lower than the 200 – 280 Bq m^−3^ measured at the surface laboratory (64). Nematodes used in BBR experiments were acclimated to this condition during two generations prior to experiments.

### Fertility assay

L4 larvae were picked individually from a synchronized population and put in separate fresh plates. Animals were transferred to fresh plates every 24 h until egg laying stopped. Synchronized animal populations were inoculated after hatching. The number of nonviable (*N*) and unfertilized eggs (*U*) was counted 24 h after the adult was removed from the plate to ensure all eggs had hatched. Viable progeny (*A*) was counted 48 h after the adult was removed from the plate. Reproductive success, *R*, was then computed as *R* = *A*/(*A* + *N* + *U*).

For OrV infection experiments, a population of animals was inoculated one hour after hatching with the OrV stock described below.

### Viral stock preparation, virus quantification and inoculation procedure

For OrV (strain JUv1580_vlc) stock preparation, SFE2 animals were inoculated as previously described (65). In short, animals were allowed to grow for 5 days and then resuspended in M9 (0.22 M KH_2_PO_4_, 0.42 M Na_2_HPO_4_, 0.85 M NaCl, 0.001 M MgSO_4_), let stand for 15 min at room temperature, vortexed, and centrifuged for 2 min at 400 g. The supernatant was centrifuged twice at 21,000 g for 5 min and then passed through a 0.2 μm filter. RNA of the resulting viral stock was extracted using the Viral RNA Isolation kit (NZYTech). The concentration of viral RNA was then determined by RT-qPCR using a standard curve, and normalized across different stocks (details below). Primers used for RT-qPCRs can be found in Table S2.

For the standard curve, cDNA of JUv1580_vlc was obtained using AccuScript High-Fidelity Reverse Transcriptase (Agilent) and reverse primers at the 3’ end of the genome. Approximately 1000 bp of the 3’ end of RNA2 were amplified using forward primers containing 20 bp coding the T7 promoter and DreamTaq DNA Polymerase (Thermo Fisher). The PCR products were gel purified using MSB Spin PCRapace (Invitek Molecular) and an *in vitro* transcription was performed using T7 Polymerase (Merck). The remaining DNA was then degraded using DNAse I (Life Technologies). RNA concentration was determined by NanoDrop (Thermo Fisher) and the number of molecules per µL was determined using the online tool EndMemo RNA Copy Number Calculator (https://www.endmemo.com/bio/dnacopynum.php). Primers used for the standard curve can be found in Table S2.

For inoculation experiments, synchronized populations were inoculated by pipetting 60 μL of viral stock on top of the bacterial lawn containing the animals. The normalized inoculum contained 2.6·10^7^ copies of OrV RNA2/μL. The efficiency of this viral stock (measured as animals showing activation of the *pals-5p::GFP* reporter at 48 hpi) was 72 ±3 % (mean ±1 SEM, *n* = 5 plates with 44 – 48 animals per plate).

### RNA extractions and RT-qPCRs

Sample preparation and RNA extractions were performed as previously described (65). Synchronized populations of 300 inoculated and control animals were collected at the designated times (from 6 hpi to 44 hpi every 2 h) with PBS-0.05% Tween. Samples of inoculated animals were performed in triplicates. Samples were centrifuged for 2 min at 1350 rpm and the supernatant was discarded. Another two wash steps were performed before flash-freezing the samples in liquid N_2_. 500 µL of Trizol (Invitrogen) were added to the nematode pellet and the pellet was disrupted by following five cycles of freeze-thawing and five cycles of 30 seconds of vortex followed by 30 seconds of rest. 100 µL of chloroform were then added and the tubes were shaken for 15 s and let rest for 2 min. Samples were centrifuged for 15 min at 11,000 g at 4 °C and the top layer containing the RNA was then mixed with the same volume of 100% ethanol. The sample was then loaded into RNA Clean & Concentrator columns (Zymo Research) and the rest of the protocol was followed according to manufacturer instructions.

RT-qPCRs were performed using Power SYBR Green PCR Master Mix (Applied Biosystems) on an ABI StepOne Plus Real-time PCR System (Applied Biosystems). 10 ng of total RNA were loaded and a comparison with the endogenous gene *cdc-42* was used for relative OrV quantifications. Primers used for RT-qPCRs can be found in Table S2.

### Body size measurements and analysis of abnormal phenotypes

Synchronized animals were collected at 12, 24, 36, and 48 h, plates were washed with M9 buffer to collect nematodes and another M9 wash was made to remove bacteria from the media. Animals were mounted in a 4% agarose pad with 20% 100 mM azide and photographed using a Leica Thunder DMi8 microscope with DFC9000 GTC sCMOS camera and objective HC PL APO 40×/0.95 CORR PH2 (Leica Microsystems). Body length was measured using the segmented line tool in ImageJ (66).

The photographs employed to measure body size were also used to describe abnormal phenotypes.

### Intestinal permeability assay

The intestinal permeability assay was performed as previously described with minor modifications (67). Synchronized nematodes were washed after 48 h with S buffer (6.5 mM K_2_HPO_4_, 43.5 mM KH_2_PO_4_, 100 mM NaCl) and suspended in a 1:1 solution of 1 mL of M9 buffer mixed with 0.05 g of erioglaucine disodium salt (Thermo Scientific) and 1 mL of *E. coli* OP50 suspended in S buffer, and gently rocked for 3 h.

Animals were washed carefully three times with 10 mL of M9 buffer. Animals were mounted on microscopic slides on top of a 4% agarose pad with 20% 100 mM azide. Photographs were taken using a Leica MZ10F stereomicroscope with Flexacam C3 camera and with a Leica DMi1 microscope with Flexacam C1 camera. Quantifications were randomized by assigning random names to photographs to avoid observer bias. A positive control was prepared to help us to classify the different phenotypes. Positive control was prepared by following the same assay except that animals were grown in NGM plates with 0.8 mM H_2_O_2_ during 48 h, as previously described (68). Four-hundred nematodes were used in each condition.

### A model of *C. elegans* fecundity and parameter estimation

Fig. 5 describes a simple fecundity model for sexually mature adults at L4 developmental stage, in which a fixed number of spermatocyte cells (*S*) have already been produced and stored in the spermathecae. Germline precursor cells (*G*) differentiate into mature oocytes (*O*) at rate *δ_G_*. This rate is assumed to be constant *δ_G_ = δ* as far as germline cells exist (*G*(*t*) > 0) and zero otherwise. These oocytes migrate from the distal to the proximal gonads at rate *η* until reaching the spermathecae where, if spermatocytes are available, are fecundated with an efficiency function *ψ* that depends on the amount of *S* present in the spermatheca at time *t*: *ψ*(*S*) = (1 − *γ*)*S*⁄(*γ* + *S*) ∈ [0,1], where *γ* is a dimensionless parameter which modulates the value of the efficiency, resulting in fertilized and unfertilized (*U*) eggs. After passing through the uterus, the eggs are deposited in the agar, where, if they have been fecundated, a fraction μ will develop into L1 larvae (*i.e*, the viable progeny, *A*). A fraction 1 − μ will stop embryonic development, resulting in nonviable embryos (*N*). *N* and *U* eggs will simply remain in the plates and are easily distinguishable.

**FIG 5.**
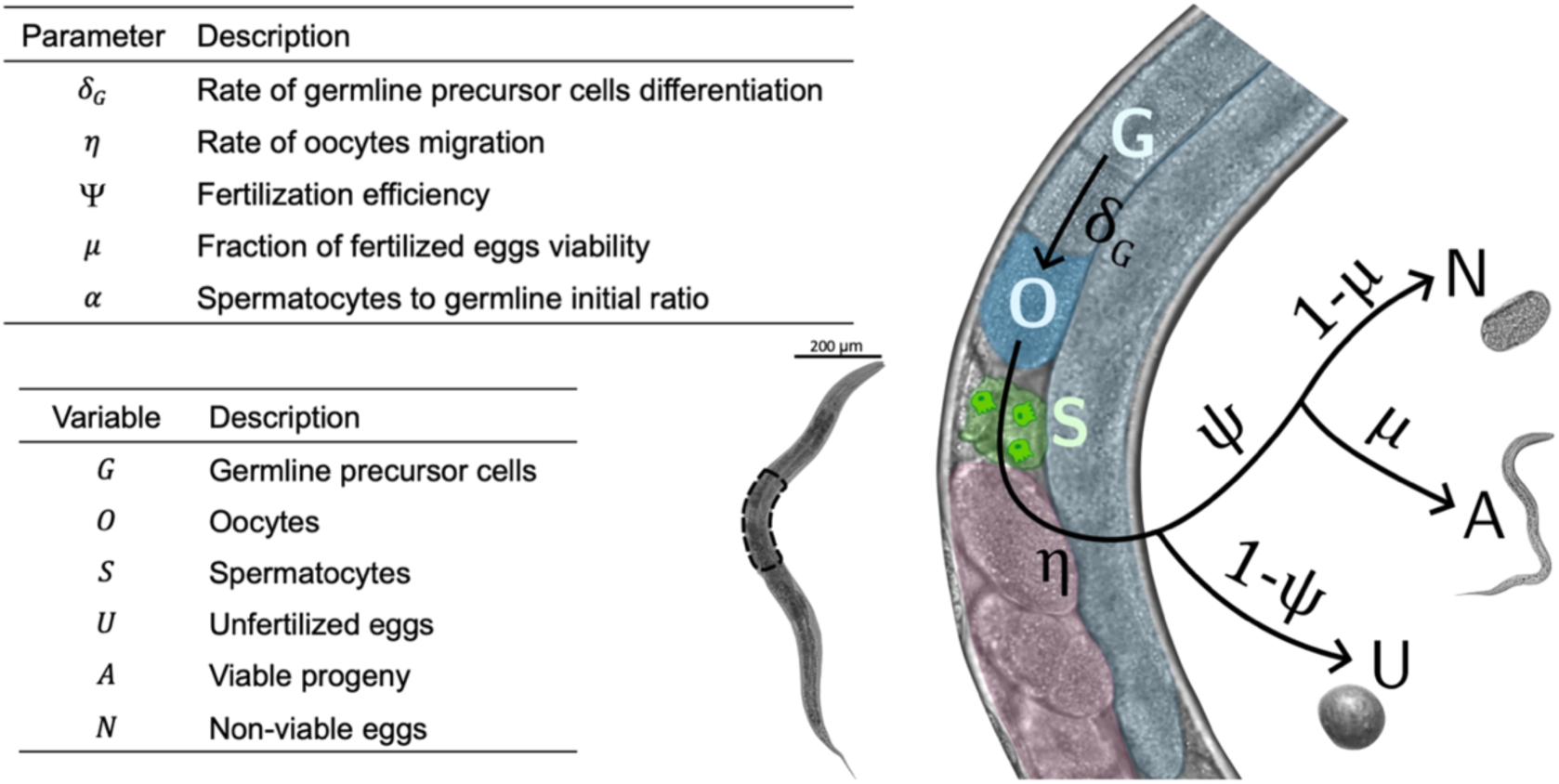
Parameters, variables and schematic representation of the fecundity model described by Eq. 1 – 6.

These processes can be written according to the following set of ordinary differential equations,

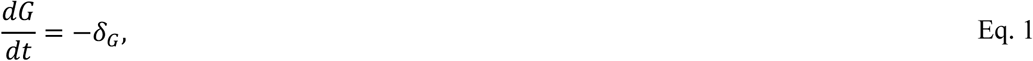

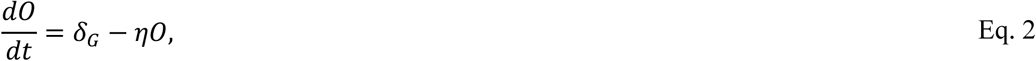

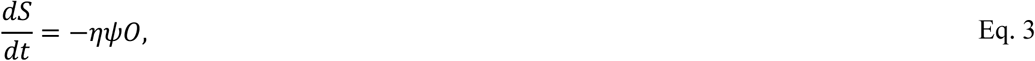

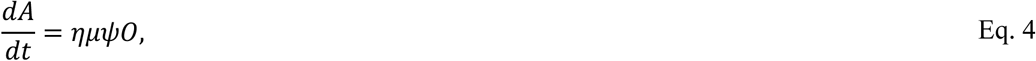

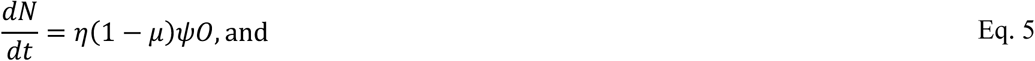

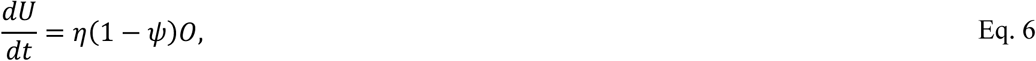

being *A*, *N*, and *U* are the three experimental observables (Eqs. 4 – 6). All variables were normalized by the initial number of non-differentiated germinal cells *G*_0_. Model parameters *δ, η* and *γ* represent prezygotic stages, while μ represents a postzygotic event. Furthermore, we define the dimensionless parameter *α* = *S*_0_/*G*_0_, which for a hermaphrodite in which the spermatocytes will be the limiting factor for asexual reproduction will be α < 1.

We used a genetic algorithm to estimate the vector of parameters *(δ, η, μ, γ, α*, *G*_0_) better fitting the experimental data for every condition. The size of the population of vectors of parameters was fixed at 600, and the number of generations was set to 10^4^. The exploration of the parameter space was refined through successive generations. The cost function (ℱ) to minimize was the logarithm of the square sum of discrepancies:

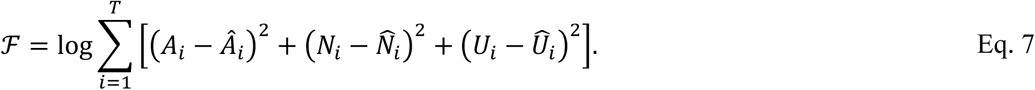

Here, 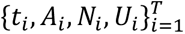 represent the experimental data obtained for *T* different times and 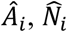 and 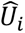 denote the estimated values for the different species obtained with the mathematical model for a given set of parameters.

Within each generation, the 5% of the parameters with the lowest ℱ values were designated as the elite population, remaining constant for the next generation. The next-best 80% underwent parameter crossover, while the remaining 15%, representing the least favorable parameters, experienced random mutations. The parameter space was subdivided into a logarithmic scale for parameters *δ, η* and *γ* to ensure a more balanced exploration. The model was fitted to every fertility assay data, resulting in a vector of estimated parameters per experimental replicate and condition.

### Statistical analyses

Data were fitted to generalized linear mixed models (GLMM) with gravity intensity (standard *vs* μG), radiation level (standard *vs* BBR) and infection status (non-inoculated *vs* inoculated) incorporated as fixed orthogonal factors, whereas individual animals (subjects) and days-post inoculation (repeated measures per subject) were both modeled as random factors. In the case of counts (*A*, *N* and *U*), a Poisson distribution with a log-link function was used; in the case of *R* and body size, a Normal distribution with identity-link function was used.

Intestinal permeability and the frequency of males per population data were fitted to logistic regression models using generalized linear models (GLM) with gravity intensity and radiation level incorporated as fixed orthogonal factors and assuming a Binomial distribution and a logit-link function.

Viral load data were fitted to a GLMM with gravity intensity and radiation level incorporated as fixed orthogonal factors, whereas replicate plates (subjects) and hpi (repeated measures per subject) were both modeled as random factors, a Gamma distribution with a log-link function was used.

Estimated parameters of the model described by Eqs. 1 – 6 were independently fitted to ANOVA models with gravity intensity, radiation level and infection status incorporated as fixed orthogonal factors. Errors were assumed to be Normal.

In all cases, pairwise *post hoc* comparisons were done with the sequential Bonferroni method. The magnitude of the effects was calculated using the 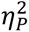 statistic, *i.e*., the proportion of variance in the dependent variable that can be attributed to each model factor after accounting for the variance explained by other factors in the model. Typically, values of 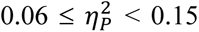 are considered as medium size effects and 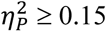 are as large magnitude effects.

These analyses were done with SPSS version 29.0.2.0 (IBM Corporation, Armonk, NY).

## ACKNOWLEDGEMENTS

We thank Francisca de la Iglesia (I2SysBio) and Rebecca Hernández-Antolín (LSC) for excellent technical support. We are deeply grateful to Carlos Peña-Garay (LSC) for his commitment and dedication to research in subterranean biology, as well as his constant material and emotional support. Many computations were performed on the HPC cluster Garnatxa at I2SysBio (CSIC-UV).

## AUTHOR CONTRIBUTIONS

A.V.G.: conceptualization, data curation, formal analysis, investigation, methodology, validation, writing-original draft, writing-review and editing; V.G.C.: conceptualization, investigation, methodology, supervision, validation, writing-review and editing; J.C.M.S.: conceptualization, formal analysis, investigation, methodology, software, writing-review and editing; E.G.L.: investigation, writing-review and editing; R.G.: conceptualization, supervision, writing-review and editing; S.F.E.: conceptualization, data curation, formal analysis, funding acquisition, project administration, supervision, writing-original draft, writing-review and editing.

All authors gave final approval for publication and agreed to be held accountable for the work performed therein.

## DATA AVAILABILITY

The datasets generated during the current study are available in the Zenodo repository, https://doi.org/10.5281/zenodo.13869555.

## ADDITIONAL FILES

Supplementary materials are available online at https://git.csic.es/sfelenalab/spaceworms/-/blob/070697e9dc66061d671094f6906a4a18f25b5e6b/Supplementary_Materials.pdf.

