## Supplementary materials for "Reduced gravity and muon flux absence affect *Caenorhabditis elegans* life history traits and viral infection"

**Text S1**

**Confirming the effect of microgravity and below-background muon radiation flux in C. elegans fecundity.**

Focusing first on the effect of both spaceflight-related stresses in the absence of infection, we found that the response of the four fecundity traits was largely different for each stress. In the case of progeny reaching adulthood (*A*), only differences in gravity intensity had a highly significant although of small magnitude effect (Table 1; *P* < 0.001, $\eta_{P}^{2}$ = 0.037). In standard gravity conditions, the grand mean *A* was 247 ±4 (±1 SE), while it was reduced to 223 ±3 in μG conditions. The effect of muon radiation flux was dependent on gravity intensity (Table 1; *P* < 0.001, $\eta_{P}^{2}$ = 0.034). While in surface radiation conditions differences in gravity intensity had no significant effects (*post hoc* test, *P* = 0.592), differences in gravity intensity had a remarkably large effect on the number of viable progeny in BBR conditions (258 ±5 in standard gravity *vs* 212 ±5 in μG; *post hoc* test, *P* < 0.001)

Neither differences in gravity intensity nor muon flux radiation had a measurable effect on the average number of unfertilized eggs (Table 1; in all cases *P* ≥ 0.166, $\eta_{P}^{2}$ ≤ 0.003).

In the case of the number of nonviable embryos (*N*), the reduction in the muon radiation flux had a significant positive effect, yet of small magnitude (Table 1; *P* = 0.026, $\eta_{P}^{2}$ = 0.008), being the grand mean *N* in standard conditions 3.2 ±0.2 and increasing to 3.9 ±0.2 in BBR conditions. Differences in gravity only had a significant effect mediated by the muon radiation flux conditions, yet of very small effect (Table 1; *P* < 0.001, $\eta_{P}^{2}$ = 0.037): in BBR conditions, the effect of μG was to reduce *N* from 4.3 ±0.3 to 3.5 ±0.3 (*post hoc* test, *P* = 0.036). In sharp contrast, in standard muon radiation flux conditions, the effect of μG was the opposite, to increase *N* from 2.1 ±0.3 to 4.3 ±0.3 (*post hoc* test, *P* < 0.001).

In conclusion from this section, both spaceflight stress conditions impacted fecundity, though their effects varied across traits. μG reduced the number of progeny reaching adulthood and reproductive success, with a more pronounced effect under BBR conditions, while having no effect on unfertilized eggs. BBR increased nonviable embryos and reduced reproductive success in standard gravity, but its effects were smaller or absent in μG. Overall, the combination of microgravity and BBR intensified the negative impacts on fecundity traits.

**OrV virulence varies with gravity and radiation conditions**

After stablishing the effect that μG and BBR had in the animal’s fecundity, we now sought to explore the effect of these stresses into the susceptibility to OrV infection (red lines and symbols in Fig. 1). Table 1 summarizes the GLMM tests on infection and its interaction with muon radiation flux levels and the intensity of gravity. Except for *U*, infection had a significant effect, by itself or in combination with spaceflight stresses, on the fecundity traits (Table 1; in all cases *P* ≤ 0.002, 0.015 ≤ $\eta_{P}^{2}$ ≤ 0.074). In the case of *A*, the grand mean viable progeny for non-inoculated animals was 224 ±4, while it was significantly larger (246 ±3) in inoculated animals (Table 1; *P* < 0.001, $\eta_{P}^{2}$ = 0.031). Indeed, the effect of infection depended on the intensity of gravity, with non-inoculated grown in standard conditions having an average progeny of 227 ±5 while those reared in μG leaving a more abundant progeny (268 ±5; *post hoc* test, *P* < 0.001). The interaction between both spaceflight-related stresses also affected the severity of infection (Table 1; *P* = 0.010, $\eta_{P}^{2}$ = 0.011). Inoculated animals grown in standard gravity and BBR conditions leaved an average progeny of 275 ±6, while the smallest progeny was observed for non-inoculated worms reared in microgravity and BBR (*i.e*., closer to real spaceflight conditions), 202 ±6.

Infection *per se* increased *N* from 3.1 ±0.2 to 4.0 ±0.2 (Table 1; *P* = 0.002, $\eta_{P}^{2}$ = 0.015), with the effect depending both on the intensity of gravity (Table 1; *P* < 0.001, $\eta_{P}^{2}$ = 0.021) and muon radiation flux (Table 1; *P* < 0.001, $\eta_{P}^{2}$ = 0.025), but not on the interaction between the two environmental factors (Table 1; *P* = 0.611, $\eta_{P}^{2}$ < 0.001). More precisely, non-inoculated animals generated the same number of nonviable embryos at both gravity intensities (3.3 ±0.3 in standard gravity *vs* 2.8 ±0.3 in μG; *post hoc* test, *P* = 0.294) while inoculated animals generated more nonviable embryos in μG conditions (from 3.2 ±0.3 to 4.9 ± 0.3; *post host* test, *P* < 0.001). The effect of differences in muon radiation flux went in the opposite direction: while non-inoculated animals generated more nonviable embryos in BBR conditions than in the surface (4.0 ±0.3 *vs* 2.1 ±0.4; *post hoc* test, *P* < 0.001), no differences were found for inoculated animals (3.8 ±0.3 *vs* 4.3 ±0.3; *post hoc* test, *P* = 0.207).

In summary from this section, OrV infection negatively impacts reproductive success, especially by reducing the number of viable progeny and increasing nonviable embryos. These effects are worsened by μG but somewhat buffered by BBR, indicating that these stresses likely affect different pathways of the infection response.

**Supplementary Figures**

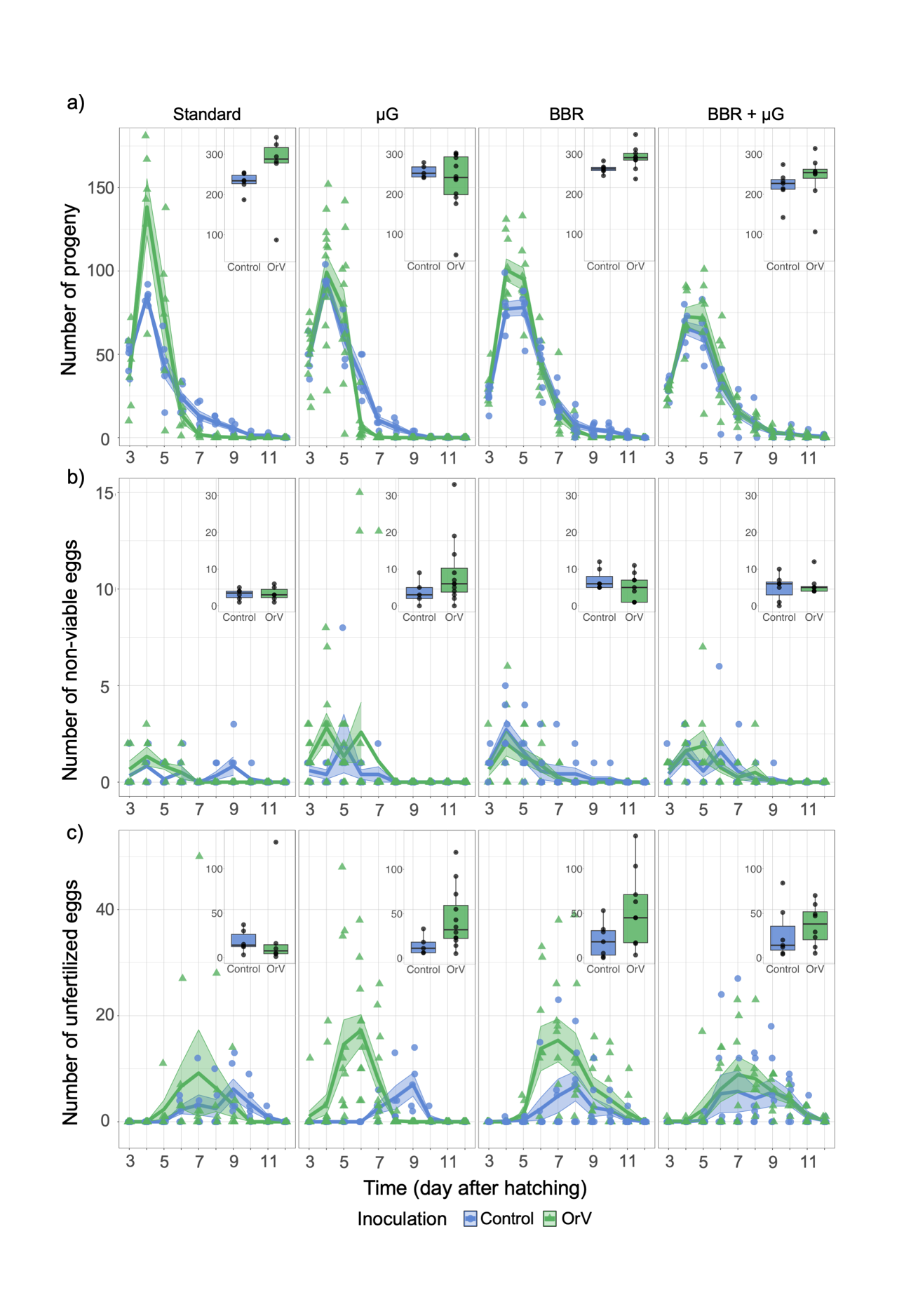

**Figure S1**. Fertility variables of OrV inoculated (green) and non-inoculated (blue) worms under control, microgravity, BBR and BBR plus microgravity conditions. Color triangles and circles represent one replicate and solid lines the mean of replicates with SE plotted as a ribbon. Boxplots in the insets represent the sum of each replicate across all day for each parameter analyzed, each dot represents a replicate. Each replicate consists of an individual worm whose egg-laying pattern has been monitored from L4 to end of progeny production. Thus, the number of replicates is consistent between parameters of fertility. For non-inoculated worms: *n* = 6 for control conditions, *n* = 5 for μG, *n* = 7 for BBR, and *n* = 7 for BBR plus μG. For OrV inoculated worms: *n* = 6 for standard conditions, *n* = 12 for microgravity, *n* = 9 for BBR, and *n* = 8 for BBR plus μG. **(a)** Number of progeny of non-inoculated and inoculated worms during the reproductive period and total of progeny. **(b)** Number of non-viable eggs of non-inoculated and inoculated worms during the reproductive period and total of non-viable eggs. **(c)** Number of unfertilized eggs of non-inoculated and inoculated worms during the reproductive period and total of unfertilized eggs. In all cases, the insets represent the area under the curves.

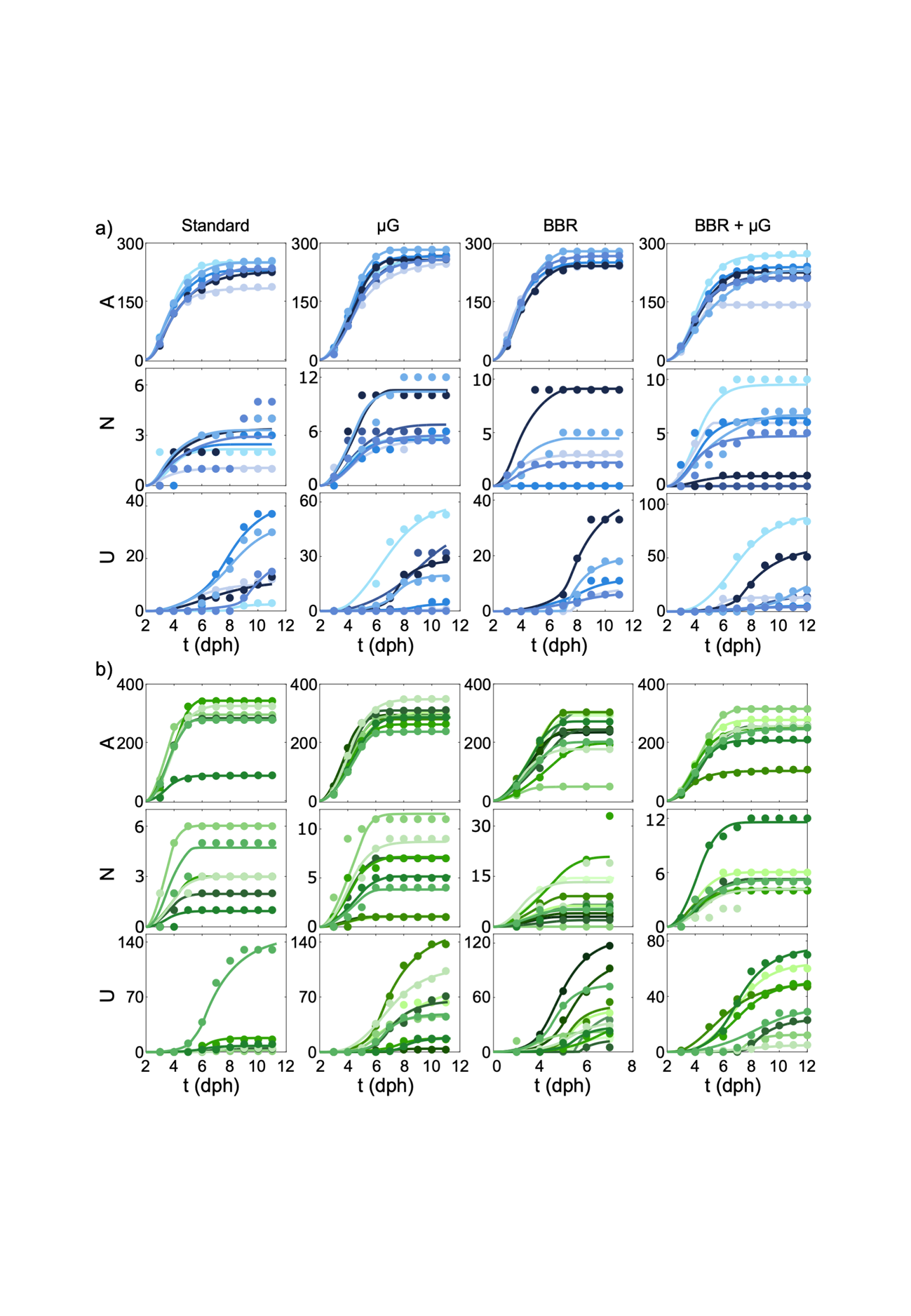

**Figure S2.** Experimental data (circles) and the corresponding fitting (solid lines) of the mathematical model using the optimized parameters’ values. *U*: unfertilized eggs; *N*: nonviable fertilized eggs; *A*: viable L1 progeny. Colors represent different biological experimental replicates. From left to right: standard conditions, BBR, μG, and combination of BBR and μG. (**a**) non-infected animals. (**b**) OrV infected animals. Parameter’s search space for the genetic algorithm: *α* [0 – 1], *δ* [10^–4^ – 10^–2^] d^–1^, *η* [10^–4^ – 10^–2^] d^–1^, *γ* [10^–4^ – 10^–1^], and *μ* [0 - 1].

| **Supplementary Tables**  **Table S1.** Summary of the 3-ways ANOVA models evaluating the effect of infection status, gravity intensity and muon radiation flux in the parameters of the fecundity model described in Eq. 1 - 6. Highlighted in dark gray are significant effects of medium or large magnitude. No significant cases were of small magnitude. | | | | | | | |
| --- | --- | --- | --- | --- | --- | --- | --- |
|  |  | ***G*_0_** | ***α*** | ***δ*** | ***η*** | ***γ*** | ***μ*** |
| Intersection | SS^a^ | 4715208.951 | 44.515 | 16.732 | 36.598 | 0.013 | 54.020 |
|  | *F*^b^ | 1021.632 | 4912.519 | 1009.616 | 374.030 | 23.412 | 206909.869 |
|  | *P* | < 0.001 | < 0.001 | < 0.001 | < 0.001 | < 0.001 | < 0.001 |
|  | $\eta_{P}^{2}$ | 0.952 | 0.990 | 0.951 | 0.878 | 0.998 | 1.000 |
| Gravity intensity | SS^a^ | 7989.144 | 0.011 | 0.025 | 6.446·10^–4^ | 0.011 | 4.519·10^–4^ |
|  | *F*^b^ | 1.731 | 1.225 | 1.507 | 0.001 | 0.803 | 1.680 |
|  | *P* | 0.194 | 0.274 | 0.225 | 0.980 | 0.374 | 0.201 |
|  | $\eta_{P}^{2}$ | 0.032 | 0.023 | 0.028 | 0.000 | 0.015 | 0.031 |
| Muon radiation flux | SS^a^ | 3690.614 | 0.009 | 0.574 | 0.733 | 4.084·10^–4^ | 0.009 |
|  | *F*^b^ | 0.800 | 0.993 | 34.626 | 7.496 | 0.726 | 0.993 |
|  | *P* | 0.375 | 0.324 | < 0.001 | 0.008 | 0.398 | 0.324 |
|  | $\eta_{P}^{2}$ | 0.133 | 0.019 | 0.400 | 0.126 | 0.014 | 0.019 |
| Infection status | SS^a^ | 20998.093 | 0.057 | 0.287 | 1.346 | 0.001 | 1.206·10^–4^ |
|  | *F*^b^ | 4.550 | 6.238 | 17.296 | 13.752 | 2.023 | 0.462 |
|  | *P* | 0.038 | 0.016 | < 0.001 | < 0.001 | 0.161 | 0.500 |
|  | $\eta_{P}^{2}$ | 0.080 | 0.107 | 0.250 | 0.209 | 0.037 | 0.009 |
| Gravity × Radiation | SS^a^ | 13387.792 | 0.003 | 0.137 | 0.045 | 3.308·10^–5^ | 2.859·10^–4^ |
|  | *F*^b^ | 2.901 | 0.328 | 8.290 | 0.461 | 0.059 | 1.095 |
|  | *P* | 0.095 | 0.569 | 0.006 | 0.500 | 0.809 | 0.300 |
|  | $\eta_{P}^{2}$ | 0.053 | 0.006 | 0.138 | 0.009 | 0.001 | 0.021 |
| Gravity × Infection | SS^a^ | 4681.164 | 0.002 | 0.013 | 0.025 | 0.002 | 0.001 |
|  | *F*^b^ | 1.014 | 0.188 | 0.797 | 0.253 | 3.233 | 2.336 |
|  | *P* | 0.319 | 0.666 | 0.376 | 0.617 | 0.078 | 0.132 |
|  | $\eta_{P}^{2}$ | 0.019 | 0.004 | 0.015 | 0.006 | 0.059 | 0.043 |
| Radiation × Infection | SS^a^ | 470.770 | 0.001 | 0.208 | 1.176 | 4.662·10^–5^ | 4.537·10^–4^ |
|  | *F*^b^ | 0.102 | 0.056 | 12.535 | 12.015 | 0.056 | 1.738 |
|  | *P* | 0.751 | 0.814 | < 0.001 | 0.001 | 0.814 | 0.193 |
|  | $\eta_{P}^{2}$ | 0.002 | 0.001 | 0.194 | 0.188 | 0.001 | 0.032 |
| Gravity × Radiation × Infection | SS^a^ | 6.008 | 0.016 | 0.039 | 0.010 | 0.001 | 1.010·10^–5^ |
|  | *F*^b^ | 0.001 | 0.001 | 2.371 | 0.097 | 1.151 | 0.039 |
|  | *P* | 0.971 | 0.971 | 0.130 | 0.757 | 0.288 | 0.845 |
|  | $\eta_{P}^{2}$ | 0.000 | 0.000 | 0.044 | 0.002 | 0.022 | 0.001 |
| Error | SS^a^ | 239999.214 | 0.471 | 0.862 | 5.088 | 0.029 | 0.014 |
| ^a^Type III sum of squares.  ^b^With 1 and 52 d.f. | | | | | | | |

| **Table S2**. Different sets of primers used and their purposes. | | |
| --- | --- | --- |
| **Primer name** | **Sequence ( 5’→3’)** | **Application** |
| oVG15_RNA2_qPCR_2_F | ACGAAGCAGTAGCCGTTAAG | RT-qPCR OrV (RNA 2) |
| oVG16_RNA2_qPCR_2_R | GAGAACATCCTTCTCTGCGG | RT-qPCR OrV (RNA 2) |
| cdc42_1_F | AGCCATTCTGGCCGCTCTCG | RT-PCR endogenous *cdc-42* gene |
| cdc42_2_R | GCAACCGCTTCTCGTTTGGC | RT-PCR endogenous *cdc-42* gene |
